# Functional overlap between the mammalian *Sar1a* and *Sar1b* paralogs in vivo

**DOI:** 10.1101/2024.02.27.582310

**Authors:** Vi T. Tang, Jie Xiang, Zhimin Chen, Joseph McCormick, Prabhodh S. Abbineni, Xiao-Wei Chen, Mark Hoenerhoff, Brian T. Emmer, Rami Khoriaty, Jiandie D. Lin, David Ginsburg

## Abstract

Proteins carrying a signal peptide and/or a transmembrane domain enter the intracellular secretory pathway at the endoplasmic reticulum (ER) and are transported to the Golgi apparatus via COPII vesicles or tubules. SAR1 initiates COPII coat assembly by recruiting other coat proteins to the ER membrane. Mammalian genomes encode two *SAR1* paralogs, *SAR1A* and *SAR1B*. While these paralogs exhibit ∼90% amino acid sequence identity, it is unknown whether they perform distinct or overlapping functions in vivo. We now report that genetic inactivation of *Sar1a* in mice results in lethality during mid-embryogenesis. We also confirm previous reports that complete deficiency of murine *Sar1b* results in perinatal lethality. In contrast, we demonstrate that deletion of *Sar1b* restricted to hepatocytes is compatible with survival, though resulting in hypocholesterolemia that can be rescued by adenovirus-mediated overexpression of either SAR1A or SAR1B. To further examine the in vivo function of these 2 paralogs, we genetically engineered mice with the *Sar1a* coding sequence replacing that of *Sar1b* at the endogenous *Sar1b* locus. Mice homozygous for this allele survive to adulthood and are phenotypically normal, demonstrating complete or near-complete overlap in function between the two SAR1 protein paralogs in mice. These data also suggest upregulation of *SAR1A* gene expression as a potential approach for the treatment of SAR1B deficiency (chylomicron retention disease) in humans.

## Introduction

The intracellular secretory pathway transports proteins from the endoplasmic reticulum (ER) to the Golgi for subsequent secretion into the extracellular space, insertion into the plasma membrane, or transport to various intracellular organelles including lysosomes and endosomes (1). Following proper folding in the ER, cargo proteins are transported to the Golgi apparatus via coat protein complex II (COPII) vesicles/tubules (2, 3). The COPII coat is a highly conserved structure consisting of five core proteins that are present in all eukaryotes: SAR1, SEC23, SEC24, SEC13, and SEC31 (4, 5). GDP-bound SAR1 is recruited to the ER exit site (ERES) by membrane-bound SEC12 to initiate coat assembly. SEC12 also functions as the guanine exchange factor for GDP-SAR1. GTP-bound SAR1 subsequently recruits the SEC23-SEC24 heterodimer complex to the ERES on the cytoplasmic face of the ER membrane via direct interaction between SAR1 and SEC23. SEC23 also acts as the guanine activating factor for SAR1, resulting in GTP hydrolysis. Following cargo recruitment and concentration in the ER lumen mediated by SEC24, SEC13 and SEC31 are recruited to form the outer coat and complete coat assembly (6).

Mammalian genomes contain multiple COPII paralogs as a result of gene duplication events, including two paralogs each for SAR1, SEC23, and SEC31, and four paralogs for SEC24 (6). Gene duplications are common evolutionary events contributing to genome expansion. One of the duplicated copies typically accumulates inactivating mutations over evolutionary time, transitioning to a pseudogene, which is eventually lost from the genome. Occasionally, one duplicate copy undergoes neofunctionalization (acquisition of new function(s) distinct from those of the ancestral gene) or subfunctionalization (division of the ancestral gene’s functions among the paralogs), resulting in maintenance of both copies in the genome. Subfunctionalization can occur at either the level of protein function or transcription (7). The expansion of COPII paralogs through the course of eukaryotic evolution is thought to have accommodated increasing secretory demand. Humans and mice deficient in one paralog typically exhibit distinct phenotypes from those deficient in the other (6, 8). In mice, loss of SEC23A results in lethality due to a neural tube closure defect whereas SEC23B deficiency leads to perinatal death from pancreatic degeneration (9, 10). However, substitution of *Sec23a* coding sequence at the *Sec23b* locus completely rescues the perinatal lethality exhibited by *Sec23b* null mice (11), demonstrating a high degree of functional overlap between these two paralogs. Mice with complete loss of SEC24A exhibit normal growth and fertility with marked hypocholesterolemia whereas mice that lack either SEC24B, SEC24C, or SEC24D exhibit lethality at various time points during development (12-15). Both SEC24A/B and SEC24C/D exhibit at least a partial degree of functional overlap (12, 16).

The SAR1A and SAR1B paralogs differ by only 20 out of 198 amino acids in humans and both appear to be expressed in a wide range of human and mouse tissues (**Figure 1**) (17, 18). Mutations in *SAR1B* are associated with the rare autosomal recessive disorder chylomicron retention disease (CMRD, or Anderson’s disease) (19), whereas a disorder due to human mutations in *SAR1A* has not been reported. Melville and colleagues previously identified three amino acid clusters differing between human SAR1A and SAR1B that alter GTPase kinetics and SEC23 affinity (20), with a recent report further demonstrating that only human SAR1A is inhibited by the alarmone ppGpp (21). While these findings suggest unique biochemical functions for these two paralogs at the molecular level, there is also evidence for some overlap in function. Genetic deletion of either *SAR1A* or *SAR1B* in Caco-2/15 cells results in decreased chylomicron secretion with a less dramatic phenotype observed in SAR1A null cells. Combined inactivation of both paralogs in Caco2/15 cells is required to recapitulate the severe phenotype seen in cells from patients with CMRD (22, 23). Similarly, while disruption of *SAR1A* is tolerated in human retinal pigment epithelial (RPE-1) cells, additional attempts to delete *SAR1B* in *SAR1A*-null cells were unsuccessful, suggesting some degree of functional compensation between these two paralogs (24).

**Figure 1.**
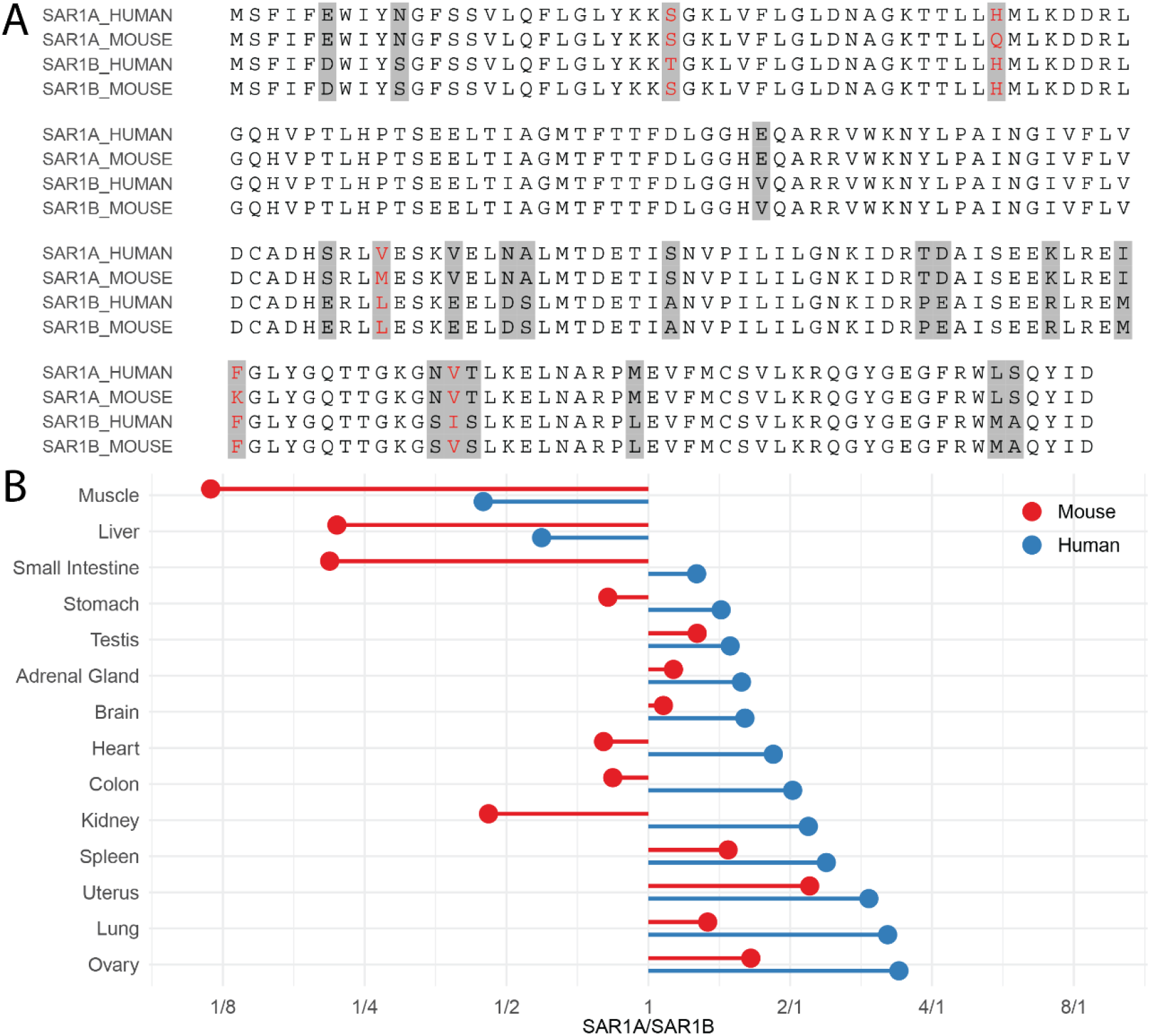
SAR1A and SAR1B are highly conserved proteins ubiquitously expressed but at variable levels across mouse and human tissues. **(A)** Protein sequence alignment of human and mouse SAR1A (NP_001136120.1 and NP_001345416.1) and SAR1B (NP_001028675.1 and NP_079811.1). Amino acid residues that differ between SAR1A and SAR1B are highlighted in grey and residues that are different between mouse and human are colored in red. **(B)** Relative *SAR1A* and *SAR1B* mRNA expression in multiple human and mouse tissues. The x-axis represents SAR1A/SAR1B mRNA abundance ratio in each tissue. Raw RNA-sequencing data were obtained from (17, 18).

In mice, germline deletion of *Sar1b* results in late-gestational lethality in homozygous null fetuses, with haploinsufficient animals demonstrating a defect in chylomicron secretion similar to that observed in homozygous CMRD patients (25). Liver specific deletion of *Sar1b* results in severe hypocholesterolemia due to a lipoprotein secretion defect (26). To date, no mouse models for SAR1A deficiency have been reported. We now report analysis of mice engineered to be genetically deficient in *Sar1a* or *Sar1b* as well as mice carrying a modified *Sar1b* locus at which the *Sar1a* coding sequence has been substituted for *Sar1b*. Our results demonstrate a high degree of functional overlap between the SAR1A and SAR1B paralogous proteins.

## Results

### Germline deletion of *Sar1a* in mice results in mid-embryonic lethality

To investigate SAR1A function in vivo, we generated mice carrying a *Sar1a* targeted allele in which *Sar1a* expression is disrupted by insertion of a gene trap cassette into intron 5 (27) (**Figure 2A**). We designed a three-primer PCR assay to differentiate between the wild type and gene trap alleles (**Figure 2B**), with DNA sequence analysis confirming the expected configuration. Quantitative PCR (qPCR) analysis of liver cDNA generated from *Sar1a*^*+/+*^ and *Sar1a*^*gt/+*^ littermates demonstrated an ∼50% reduction of *Sar1a* transcripts in the latter, whereas *Sar1b* mRNA abundance remained unchanged, consistent with inactivation of a single *Sar1a* allele (**Figure 2C**). To generate mice with homozygous loss of *Sar1a*, we performed an intercross between *Sar1a*^*gt/+*^ mice, with genotyping results shown in **Table 1**. No *Sar1a*^*gt/gt*^ mice were identified among the 65 offspring genotyped at postnatal day 21 (P21). The results of genotyping for embryos harvested at embryonic days E10.5, E11, and E11.5 post coitus (pc) are shown in **Table 1**. Though the expected number of *Sar1a*^*gt/gt*^ embryos were observed at E10.5, no viable *Sar1a*^*gt/gt*^ embryos remained by E11.5. Histologic evaluation of E10.5 embryos failed to identify any obvious abnormalities in the *Sar1a*^*gt/gt*^ embryos.

**Table 1:** Germline loss of SAR1A results in lethality during mid-embryogenesis.

| Mating<br>Genotype<br>Expected | *Sar1a<sup>gt/+</sup>* X *Sar1a<sup>gt/+</sup>* | | | p-value ($\chi^2$<br>test) |
| --- | --- | --- | --- | --- |
|  | *Sar1a<sup>+/+</sup>*<br>25% | *Sar1a<sup>gt/+</sup>*<br>50% | *Sar1a<sup>gt/gt</sup>*<br>25% |  |
| E10.5 | 10 (28%) | 16 (44%) | 10 (28%) | 0.800 |
| E11.0 | 5 (26%) | 10 (53%) | 4 (21%) | 0.924 |
| E11.5 | 7 (32%) | 15 (68%) | 0* (0%) | 0.025 |
| Weaning (P21) | 17 (26%) | 48 (74%) | 0 (0%) | 7.22 x 10<sup>-6</sup> |

\*Two *Sar1a<sup>gt/gt</sup>* embryos appeared dead and were partially resorbed.

**Figure 2.**
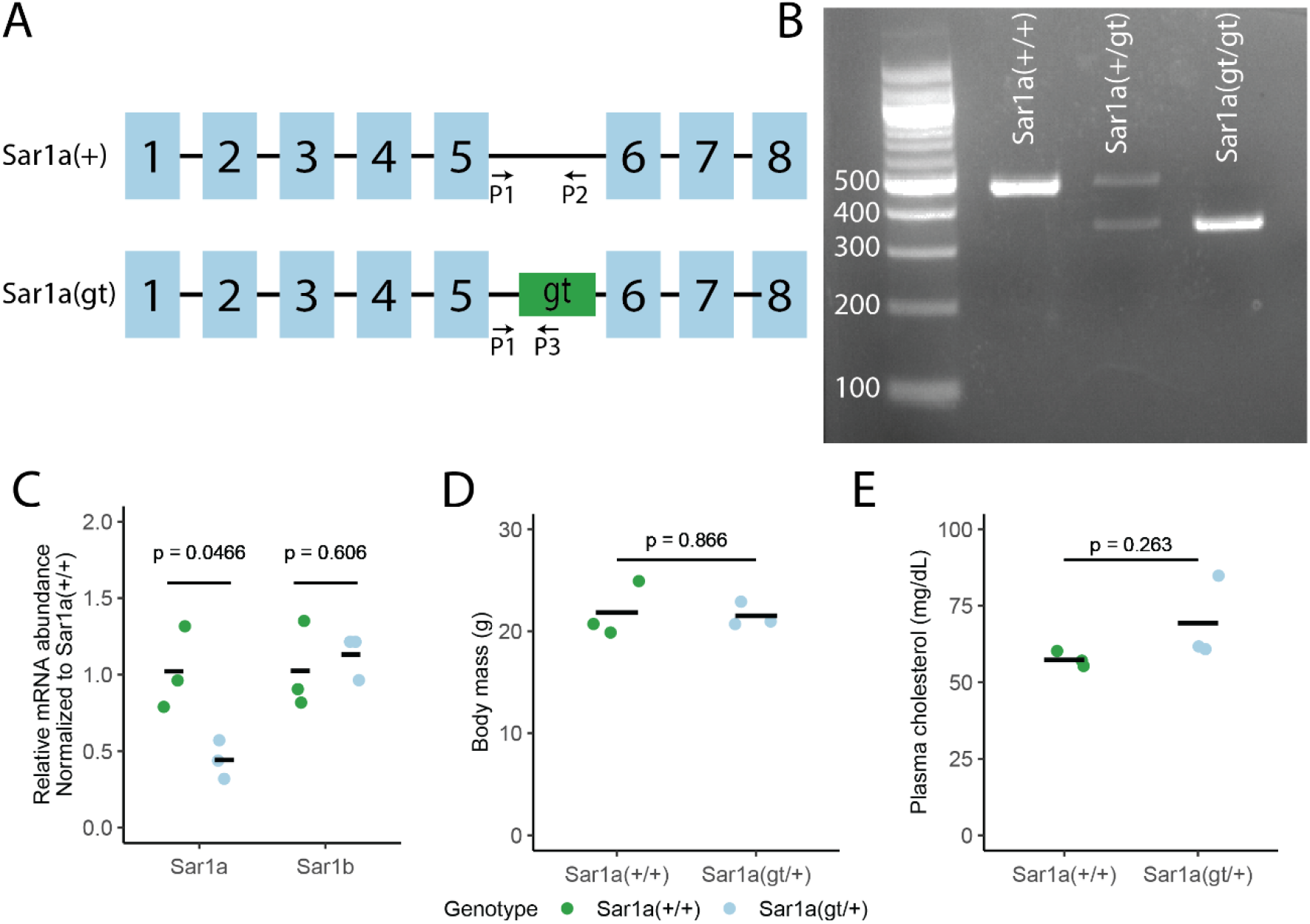
Haploinsufficiency of SAR1A is tolerated in mice. **(A)** Schematic of the wild type *Sar1a(+)* and gene trap *Sar1a(gt)* alleles The gene trap cassette is inserted into intron 5 of the *Sar1a* gene. P1, P2, P3 denote the binding sites of genotyping primers. Rectangles represent exons and solid line segments represent introns. Exons and introns are not drawn to scale. **(B)** Genotyping assay using the primers denoted in (A) and genomic DNA isolated from embryonic yolk sacs. The *Sar1a(+)* allele produces a PCR amplicon of 489 bp whereas the *Sar1a(gt)* allele produces an amplicon of 352 bp. **(C)** Liver *Sar1a* and *Sar1b* mRNA abundance in *Sar1a*^*gt/+*^ and littermate controls determined by qPCR (normalized to the level of *Sar1a*^*+/+*^ samples). **(D-E)** Body mass and plasma cholesterol levels of wild type and heterozygous *Sar1a*^gt/+^ mice. For panels (C-E) Statistical significance was determined by two-sided t-test.

Heterozygous *Sar1a*^*gt/+*^ mice were present at the expected Mendelian ratio at weaning (P21) and exhibit normal growth and fertility. At 8 weeks of age, no significant differences in body mass or plasma cholesterol levels were observed between *Sar1a*^*+/+*^ and *Sar1a*^*gt/+*^ mice (**Figure 2D-E**).

### Germline homozygous *Sar1b* deletion results in perinatal lethality

We generated heterozygous *Sar1b* deficient mice (*Sar1b*^*+/-*^) by crossing mice carrying a conditional *Sar1b* allele (*Sar1b*^*fl/+*^) with mice carrying a ubiquitously expressed Cre (*EIIa-Cre*) transgene. Cre-mediated excision of *Sar1b* exon 5 (**Figure 3A**) was confirmed with a three-primer PCR assay clearly differentiating the wild type and null alleles (**Figure 3B**). As shown in **Figure 3C**, *Sar1b* mRNA level in livers collected from *Sar1b*^*+/-*^ mice was reduced to approximately 50% of that in *Sar1b*^*+/+*^ littermates, with no difference in *Sar1a* transcript abundance between the two groups. The results of a heterozygous *Sar1b*^*+/-*^ intercross are shown in **Table 2**. Though no homozygous *Sar1b*^*-/-*^ mice were observed at weaning, the expected numbers of *Sar1b*^*-/-*^ embryos were present at E11.5 and E18.5, with most *Sar1b*^*-/-*^ neonates appearing to die shortly after birth (**Table 2**). Histologic analysis of E18.5 embryos failed to identify any obvious abnormalities in *Sar1b*^*-/-*^ embryos.

**Table 2:** Genetic inactivation of *Sar1b* leads to perinatal lethality.

| Mating<br>Genotype<br>Expected | *Sar1b<sup>+/-</sup>* X *Sar1b<sup>+/-</sup>* | | | p-value ($\chi^2$<br>test) |
| --- | --- | --- | --- | --- |
|  | *Sar1b<sup>+/+</sup>*<br>25% | *Sar1b<sup>+/-</sup>*<br>50% | *Sar1b<sup>-/-</sup>*<br>25% |  |
| E11.5 | 3 (19%) | 8 (50%) | 4 (31%) | 0.779 |
| E18.5 | 14 (26%) | 24 (44%) | 16 (30%) | 0.665 |
| P0 | 4 (31%) | 7 (54%) | 2 (15%) | 0.707 |
| P1 | 4 (57%) | 3 (43%) | 0 (0%) | 0.095 |
| Weaning (P21) | 29 (35%) | 54 (65%) | 0 (0%) | 9.21 x 10<sup>-7</sup> |

**Figure 3.**
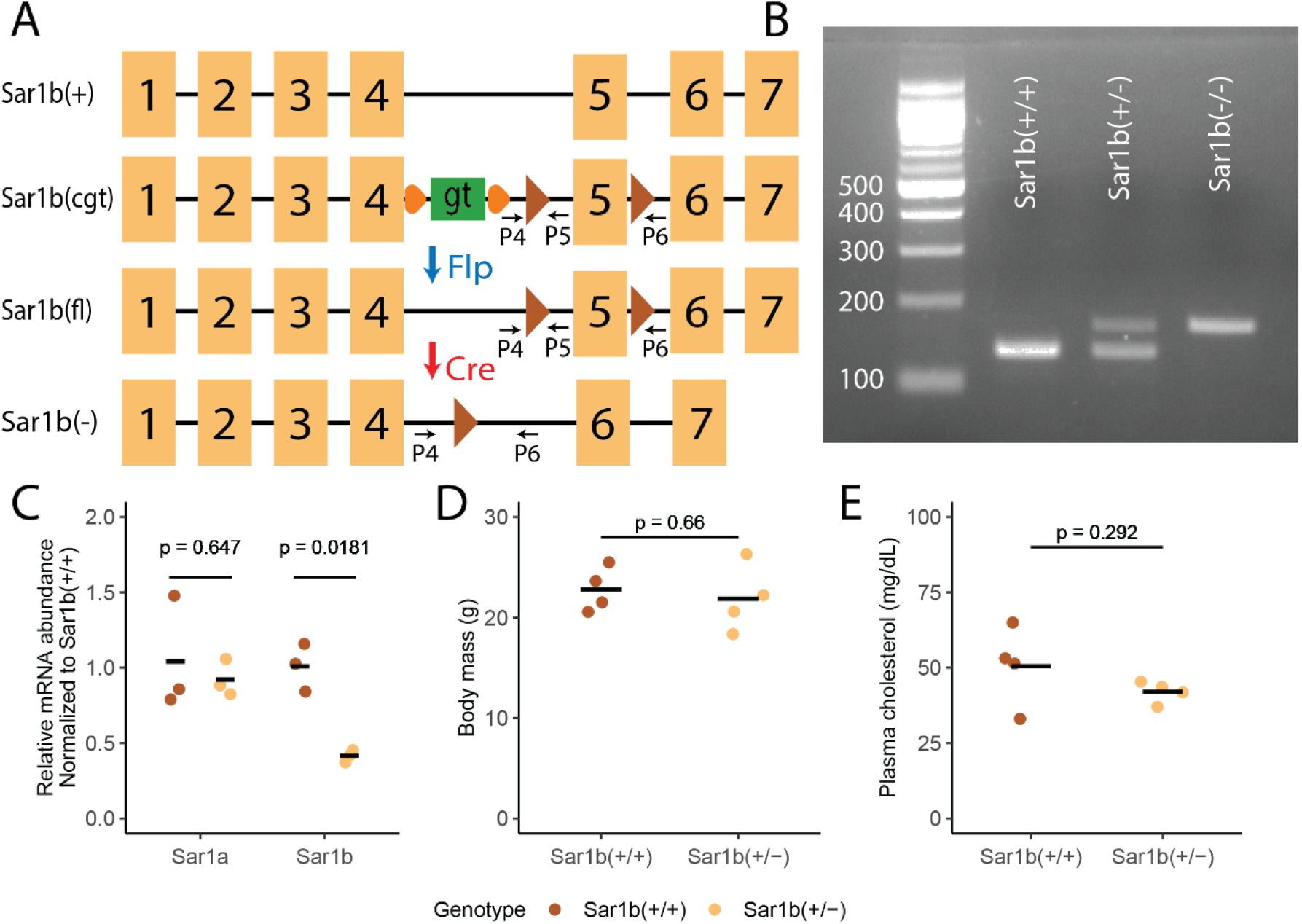
Mice with heterozygous loss of *Sar1b* demonstrate normal growth and plasma cholesterol levels. **(A)** Schematic of the wild-type *Sar1b(+)*, condition gene trap *Sar1b(cgt)*, conditional *Sar1b(fl)* and the null *Sar1b(-)* alleles. The gene trap (gt) cassette is inserted into intron 4 of the *Sar1b* gene. Orange circles represent the two FRT sites flanking the gene trap cassette. The Flp recombinase excises the gene trap cassette, resulting in the *Sar1b(fl)* allele. In the *Sar1b(fl)* allele, exon 5 of the *Sar1b* gene is flanked by two loxP sites (brown triangles), which undergo recombination in the presence of Cre recombinase, leading to excision of exon 5 in the *Sar1b(-)* allele. P4, P5, P6 denote the binding sites of genotyping primers. Rectangles represent exons and solid line segments represent introns. Exons and introns are not drawn to scale. **(B)** Genotyping assay using primers denoted in (A) and genomic DNA isolated from tail clips. The *Sar1b(+)* allele generates a PCR product of 136 bp whereas the *Sar1b(-)* allele produces a PCR amplicon of 169 bp. **(C)** Relative liver *Sar1a* and *Sar1b* mRNA abundance in *Sar1b*^*+/-*^ mice and littermate controls determined by qPCR (normalized to the mean level of *Sar1b*^*+/+*^ samples). **(D-E)** Body mass and plasma cholesterol levels of wild type and heterozygous *Sar1b*^+/-^ mice. For panels (C-E), statistical significance was determined by two-sided t-test.

Heterozygous *Sar1b*^*+/-*^ mice were observed at the expected numbers at weaning and exhibited normal growth and fertility (**Table 2**). At 3-4 months of age, no differences in body mass or plasma cholesterol levels were observed between *Sar1b*^*+/-*^ and *Sar1b*^*+/+*^ mice (**Figure 3D-E**).

### Hypocholesterolemia resulting from hepatic SAR1B deficiency is rescued by overexpression of either SAR1A or SAR1B

We next inactivated *Sar1b* in hepatocytes by combining an *Alb-Cre* transgene and the *Sar1b*^*fl*^ allele. *Sar1b*^*fl/fl*^ *Alb-Cre*^*+*^ mice were viable and exhibited a marked reduction in plasma cholesterol levels to approximately 20 mg/dL (compared to ∼75 mg/dL in littermate controls, **Figure 4A**). No reduction in cholesterol was observed in mice with hepatic haploinsufficiency for *Sar1b* (*Sar1b*^*fl/+*^ *Alb-Cre*^*+*^, **Figure 4A)**, consistent with the normal plasma cholesterol level of *Sar1b*^*+/-*^ mice (**Figure 3E**).

**Figure 4:**
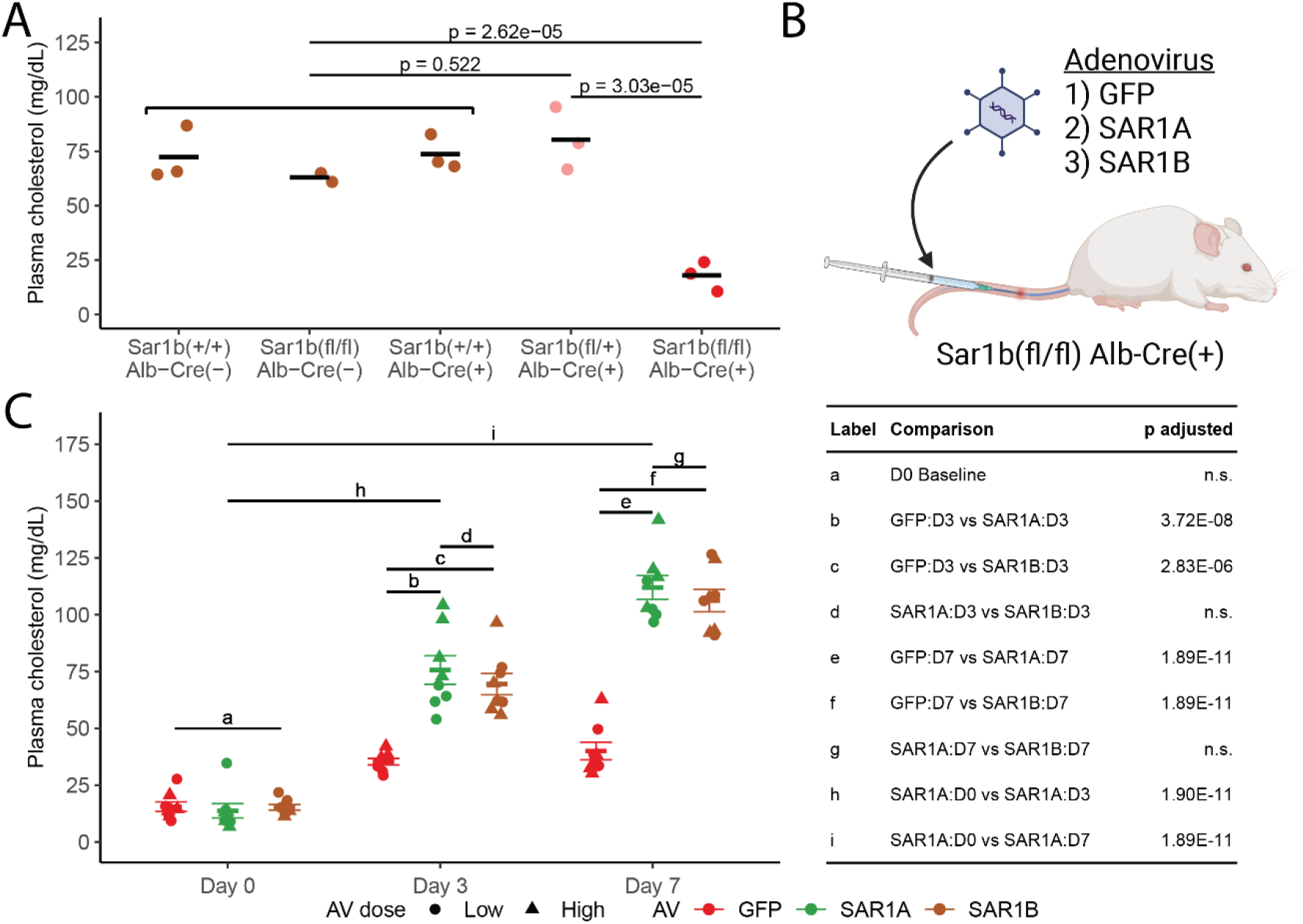
Hypocholesterolemia in hepatic SAR1B deficient mice is rescued by overexpression of either SAR1A or SAR1B. **(A)** Plasma cholesterol levels in controls (*Sar1b*^*+/+*^ *Alb-Cre*^*-*^, *Sar1b*^*fl/fl*^ *Alb-Cre*^*-*^, and *Sar1b*^*+/+*^ *Alb-Cre*^*+*^ – brown), hepatic *Sar1b* haploinsufficient (*Sar1b*^*fl/+*^ *Alb-Cre*^*+*^ – pink), and hepatic *Sar1b* null (*Sar1b*^*fl/fl*^ *Alb-Cre*^*+*^ – red) mice. Statistical significance was determined by pairwise two-sided t-test followed by Bonferroni adjustment for multiple hypothesis testing. **(B)** Schematic of SAR1B deficiency rescue experiment. *Sar1b*^*fl/fl*^ *Alb-Cre*^*+*^ mice were injected with adenovirus (AV) expressing GFP, mouse SAR1A, or mouse SAR1B via the tail vein. Blood was sampled on day 0, 3, and 7 post-injection and assayed for cholesterol levels. **(C)** Plasma cholesterol levels of mice injected with AV as illustrated in (B). Two different AV doses were used (Low and High—see Methods; n = 4 per dose per AV). Statistical significance was determined by Two-way ANOVA test with interaction between AV type, time, and dose followed by Tukey’s post-hoc test. P-values for the labelled comparisons are listed in the table on the right; n.s., not significant. p-values for all comparisons are listed in **Table S1**.

To examine potential overlap in function between SAR1A and SAR1B, we next tested the capacity of an adenovirus (AV) expressing either SAR1A, SAR1B, or GFP (as a control) to rescue the hypocholesterolemia of *Sar1b*^*fl/fl*^ *Alb-Cre*^*+*^ mice. AVs were injected via the tail vein into *Sar1b*^*fl/fl*^ *Alb-Cre*^*+*^ mice (**Figure 4B**) and plasma cholesterol levels were determined on days 0, 3, and 7 post-injection. Injection of the SAR1B AV resulted in a rapid increase in plasma cholesterol to wild type levels by day 3, with minimal increase observed with the control GFP AV, demonstrating efficacy of the AV delivery and efficient expression of *Sar1b* in the targeted hepatocytes. Injection of the SAR1A AV resulted in a similar rescue of the hypocholesterolemia phenotype, indistinguishable from that observed with SAR1B AV (**Figure 4C**). These data demonstrate that SAR1A can compensate for SAR1B deficiency in this assay, suggesting significant overlap in function between the SAR1 paralogs, at least in hepatocytes.

### Expression of *Sar1a* coding sequence at the *Sar1b* endogenous locus rescues the lethal *Sar1b* null phenotype in mice

To further examine the overlap in function between SAR1A and SAR1B in vivo, we generated a *Sar1b*^*a*^ allele in which the *Sar1b* coding sequence is replaced with that of *Sar1a* at the *Sar1b* endogenous locus, beginning immediately downstream of the ATG start codon in exon 2 of *Sar1b* (**Figure 5A**). Mice homozygous for this allele should be unable to express the SAR1B protein, with SAR1A expressed under the transcriptional control of both the *Sar1a* and *Sar1b* endogenous regulatory elements. A three-primer PCR genotyping assay was developed that could clearly differentiate the wild type and *Sar1b*^*a*^ alleles (**Figure 5B**).

**Figure 5:**
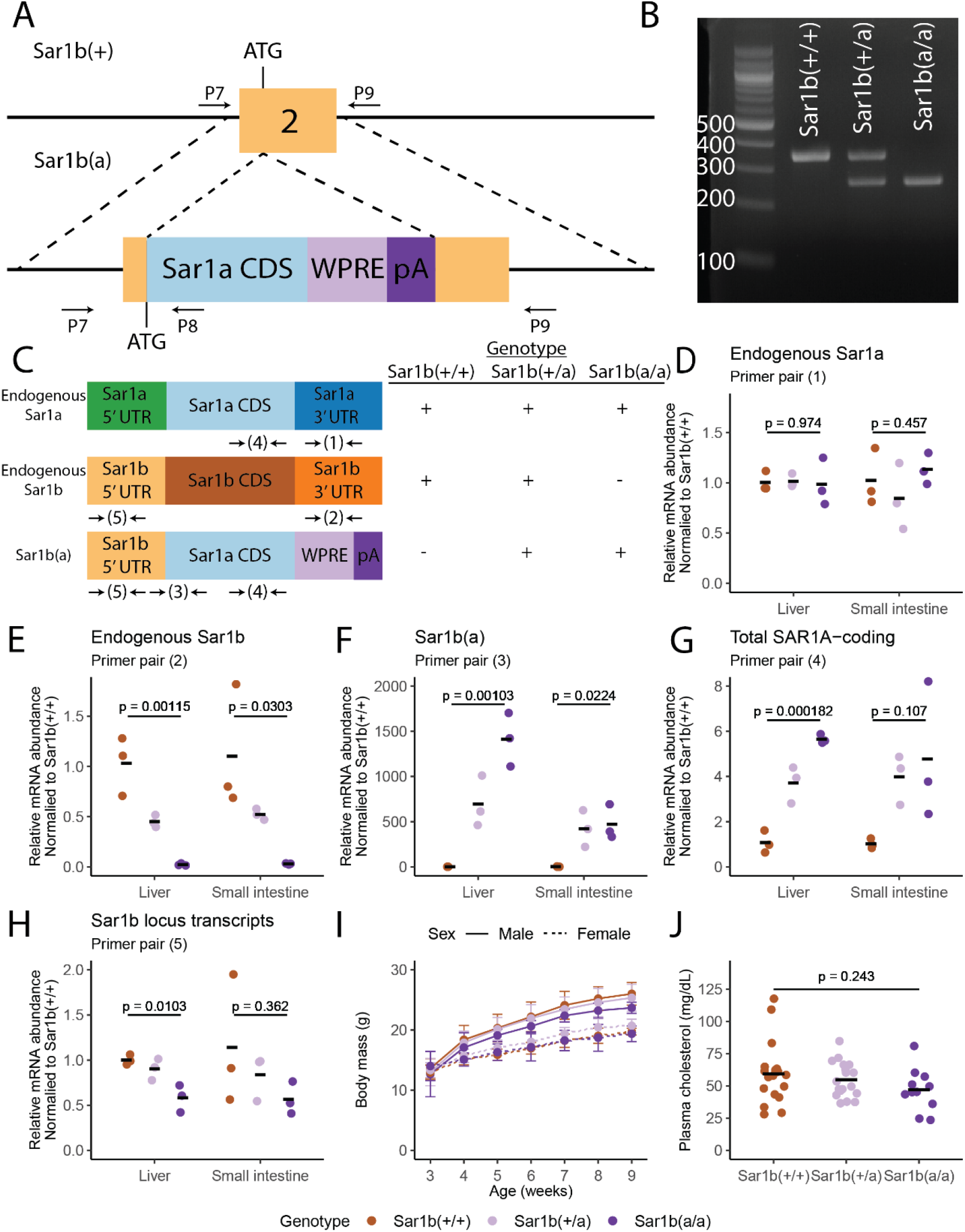
SAR1A expressed under control of the *Sar1b* endogenous locus can compensate for complete loss of SAR1B. **(A)** Schematic of the *Sar1b(a)* allele. Dashed lines mark the homology arms for DNA recombination. Exons and introns are not drawn to scale. CDS, coding sequence; WPRE, woodchuck hepatitis virus post-transcriptional regulatory element; pA, polyA sequence. P7, P8, P9 denote binding sites for genotyping primers. **(B)** Genotyping assay using primers depicted in (A) with genomic DNA isolated from tail clips. The wild type *Sar1b(+)* allele is expected to produce a PCR product of 358 bp (primer pair P7 and P9) whereas the *Sar1b(a)* allele generates a PCR amplicon of 268 bp (primer pair P7 and P8). **(C)** Schematic of the endogenous *Sar1a*, endogenous *Sar1b*, and *Sar1b(a)* transcript. The table on the right shows whether each transcript is present in the indicated genotype. Each number (with arrows on either side) below the transcript represents a particular primer pair used for detecting that transcript by qPCR. Primer pairs (1)-(3) are specific for each of the endogenous *Sar1a*, endogenous *Sar1b*, and *Sar1b(a)* transcript, respectively. Primer pair (4) amplifies both the endogenous *Sar1a* and *Sar1b(a)* transcripts (Total SAR1A-coding). Primer pair (5) amplifies both the endogenous *Sar1b* and *Sar1b(a)* transcripts (*Sar1b* locus transcript). **(D-H)** Relative abundance for transcripts as determined by qPCR using the primers illustrated in (C) (normalized to the mean level of *Sar1b*^*+/+*^ samples) in livers and small intestines collected from *Sar1b*^*+/+*^, *Sar1b*^*+/a*^, and *Sar1b*^*a/a*^ littermates. Statistical significance was determined by a one-way ANOVA test. **(I)** Body mass of male and female offspring from *Sar1b*^*+/a*^ X *Sar1b*^*+/a*^ intercrosses between 3 and 9 weeks of age (n = 23, 35, and 6 for from *Sar1b*^*+/+*^, *Sar1b*^*+/a*^, and *Sar1b*^*a/a*^, respectively; error bars represent the standard deviation for each group). **(J)** Plasma cholesterol levels of *Sar1b*^*+/+*^, *Sar1b*^*+/a*^, and *Sar1b*^*a/a*^ mice at 3-4 months of age. Statistical significance was determined by a one-way ANOVA test.

To confirm expression of SAR1A under control of *Sar1b* endogenous regulatory elements in *Sar1b*^*a/a*^ mice, we isolated RNA from the liver and small intestine of these mice and designed qPCR assays that could distinguish the endogenous *Sar1a(+)* and *Sar1b(+)* transcripts, the *Sar1b(a)* transcript, as well as primers that amplify both the *Sar1a(+)* and *Sar1b(a)* transcripts (total SAR1A-coding mRNA), or the *Sar1b(+)* and *Sar1b(a)* transcripts (total mRNA from the *Sar1b* locus) (**Figure 5C**). No significant differences were observed in the abundance of endogenous *Sar1a* mRNA between *Sar1b*^*+/+*^, *Sar1b*^*+/a*^, *Sar1b*^*a/a*^ mice (**Figure 5D**). However, *Sar1b*^*+/a*^ and *Sar1b*^*a/a*^ mice demonstrated a dose-dependent decrease in endogenous *Sar1b* and a corresponding increase in total SAR1A-coding transcript levels as compared to *Sar1b*^*+/+*^ mice (**Figure 5E and 5G**). Although *Sar1b*^*a*^ transcripts were only detected in *Sar1b*^*+/a*^ and *Sar1b*^*a/a*^ (and not *Sar1b*^*+/+*^) mice (**Figure 5F**), *Sar1b*^*a*^ mRNA expression was reduced to ∼60% of the wild-type *Sar1b* gene (**Figure 5H**).

As shown in **Table 3**, intercrosses between *Sar1b*^*+/a*^ mice generated homozygous *Sar1b*^*a/a*^ mice that are present at weaning (though at a slightly lower percentage than the expected Mendelian ratio) and demonstrate normal growth (**Figure 5I**) and fertility. Subsequent *Sar1b*^*+/a*^ X *Sar1b*^*a/a*^ crosses produced *Sar1b*^*a/a*^ offspring at the expected ratio (**Table 3**). Histologic examination of multiple organs did not identify any abnormalities in *Sar1b*^*a/a*^ mice. *Sar1b*^*a/a*^ mice also demonstrated plasma cholesterol levels indistinguishable from wild type *Sar1b*^*+/+*^ and heterozygous *Sar1b*^*+/a*^ mice (**Figure 5J**).

**Table 3:** Expression of SAR1A under the control of *Sar1b* locus rescues the lethality in *Sar1b* null mice.

| Mating<br>Genotype<br>Expected | *Sar1b<sup>+/a</sup>* X *Sar1b<sup>+/a</sup>* | | | p-value ($\chi^2$ test) |
| --- | --- | --- | --- | --- |
|  | *Sar1b<sup>+/+</sup>*<br>25% | *Sar1b<sup>+/a</sup>*<br>50% | *Sar1b<sup>a/a</sup>*<br>25% |  |
| Weaning | 80 (30%) | 148 (55%) | 43 (15%) | 0.002 |
| Mating<br>Genotype<br>Expected | *Sar1b<sup>+/a</sup>* X *Sar1b<sup>a/a</sup>* |  |  |  |
|  | *Sar1b<sup>+/a</sup>*<br>50% | *Sar1b<sup>a/a</sup>*<br>50% |  |  |
| Weaning | 23 (52%) | 21 (48%) |  | 0.763 |

## Discussion

In this study, we investigate the in vivo function of the two closely related SAR1 paralogs in mice, demonstrating a high degree of overlap. Though both paralogs are required for survival to weaning, complete deficiency for SAR1A results in lethality at mid-embryogenesis, while *Sar1b* null mice die shortly after birth. Humans with homozygous or compound heterozygous loss-of-function mutations in *SAR1A* have not been reported to date, consistent with an early embryonic lethality phenotype similar to that seen in mice. The perinatal death of *Sar1b* null mice contrasts with the human phenotype, with homozygous *SAR1B* loss-of-function mutations in humans resulting in CMRD, characterized by plasma lipid abnormality and survival to adulthood. Nonetheless, haploinsufficiency in either SAR1A or SAR1B is well tolerated in mice and human as demonstrated in this study, and by the expected prevalence of heterozygous loss-of-function mutations in either *SAR1A* or *SAR1B* among a large cohort of human subjects in the gnomAD database (28).

Despite the high degree of amino acid sequence similarity between the two SAR1 paralogs, complete loss of each paralog in mice results in death at distinct developmental time points. Taken together with previously reported differences in GTP hydrolysis and affinity for SEC23B (20), these data suggest that SAR1A and SAR1B have diverged in protein function. However, we observed that overexpression of SAR1A in the liver rescued the hypocholesterolemia phenotype of mice with hepatocyte-specific SAR1B deficiency (*Sar1b*^*fl/fl*^ *Alb-Cre*^*+*^), demonstrating at least partial overlap in SAR1A and SAR1B function in hepatocytes, although confounding effects from SAR1A overexpression or an indirect effect of AV infection cannot be excluded. Of note, there was a small but significant increase in cholesterol levels in mice treated with the control AV (expressing GFP) in this experiment. To address these issues, and to examine the potential for paralog specific function outside of hepatocytes, we generated mice expressing the SAR1A coding sequence under the transcriptional control of both its endogenous locus and the *Sar1b* locus. The complete rescue of the lethality caused by complete loss of SAR1B expression by the *Sar1b*^*a*^ allele demonstrates extensive, and potentially complete, overlap in function between the SAR1A and SAR1B proteins in vivo.

Three potential mechanisms have been proposed to explain the maintenance of paralogs such as SAR1A and SAR1B in the genome following gene duplication: neofunctionalization, subfunctionalization at the transcription level, or subfunctionalization at the protein level (7). Our findings suggest that subfunctionalization at the transcription level likely explains the maintenance of the two SAR1 paralogs in mammalian genomes. Expression of both paralogs is required, with loss of a single paralog sufficient to reduce total SAR1 below a critical level in select cell types, resulting in the unique patterns of lethality for SAR1A and SAR1B deficiency in mice and the difference in SAR1B deficient phenotypes between mice and humans.

The phenotypic differences between humans and mice with SAR1B deficiency is reminiscent of that observed for humans and mice lacking SEC23B (10, 29). Divergence in gene expression patterns across tissues within the same species and across mammalian species could explain this phenomenon for both paralog pairs. The difference in the relative expression of SAR1A and SAR1B in mice and human small intestine, with SAR1B being the dominant paralog in mouse small intestine whereas SAR1A is expressed at a higher level than SAR1B in human small intestine (**Figure 1B**), could explain the survival of humans carrying homozygous loss-of-function mutations in *SAR1B* while *Sar1b*^*-*.*/-*^ mice exhibit perinatal lethality.

As noted above human SAR1A and SAR1B have been shown to exhibit distinct biochemical properties in vitro (20, 21). Despite these differences at the molecular level, our data demonstrate that the corresponding mouse paralogs are sufficiently similar in function such that SAR1A can compensate for the loss of SAR1B in vivo. We cannot exclude greater similarity in function for the mouse versus human paralogs or subtle phenotypic differences between *Sar1b*^*+/+*^ and *Sar1b*^*a/a*^ mice. Of note, we did observe a modest loss of *Sar1b*^*a/a*^ mice in the initial *Sar1b*^*+/a*^ X *Sar1b*^*+/a*^ intercross (**Table 4**). However, a subsequent *Sar1b*^*+/a*^ X *Sar1b+*^*a/a*^ cross yielded the expected number of *Sar1b*^*a/a*^ offspring. These data could be the result of mosaicism in the founder mice, stochastic variation in gene expression between individual mice, minor differences in the genetic background of the founder mice and their progeny (C57BL/6N vs. C57BL/6J), or a passenger gene effect (30).

Lastly, upregulation of a closely related gene paralog represents a potential strategy for the treatment of select genetic diseases. For example, upregulation of γ-globin expression has been demonstrated to provide dramatic clinical improvement in patients with sickle cell disease and β-thalassemia due to genetic abnormalities in its closely related paralog, β-globin (31, 32). Our data suggest that therapeutic upregulation of one SAR1 paralog could potentially rescue deficiency of the other. Indeed, CRISPR-mediated activation of SEC23A was shown to rescue the erythroid differentiation defect exhibit by a SEC23B-deficient erythroid cell line (33). A similar strategy could potentially be leveraged to treat patients with CMRD due to SAR1B deficiency through the activation of SAR1A.

## Materials and Methods

### Animal care

All animal care and use complied with the Principles of Laboratory and Animal Care established by the National Society for Medical Research. Mice were housed in a controlled lighting (12h light/dark cycle) and temperature (22°C) environment and had free access to food (5L0D, LabDiet, St. Louis, MO) and water. All animal protocols were approved by the Institutional Animal Care and Use Committee (IACUC) of the University of Michigan (protocol number PRO00011038). Both male and female mice were used in this study.

### Generation of *Sar1a*^*gt/+*^ mice

The Bay Genomics ES cell clone CSH949 (27) was obtained from the Mutant Mouse Resource & Research Centers (MMRRC). This clone is heterozygous for the *Sar1a*^*gt*^ allele in which a gene trap cassette is inserted into *Sar1a* gene intron 5 (**Figure 2A**). ES cell culture, expansion, and microinjection to generate ES cell chimeric mice were performed as previously described (14, 15). Chimeric mice were bred with C57BL/6J (0006640, Jackson Laboratory, Bar Harbor ME) to obtain germ-line transmission. Mice carrying the *Sar1a*^*gt*^ allele were maintained by continuous backcrossing to C57BL/6J.

### Generation of *Sar1b*^*+/-*^ mice

Mice carrying the *Sar1b*^*tm1a(EUCOMM)Wtsi*^ allele (34) were obtained from the European Conditional Mouse Mutagenesis (EUCOMM) program (Wellcome Trust Sanger Institute, Cambridge, UK). These mice carry a conditional gene trap (*Sar1b*^*cgt*^) allele (**Figure 3A**) in which the gene trap cassette flanked by FRT sites followed by a loxP site is inserted into intron 4 of the *Sar1b* gene. An additional loxP site is inserted downstream of exon 5 of *Sar1b*. Mice carrying the *Sar1b*^*cgt*^ allele were maintained by continuous backcrossing to C57BL/6J mice.

To generate the conditional *Sar1b* allele (*Sar1b*^*fl*^, **Figure 3A**), *Sar1b*^*cgt/+*^ mice were crossed with mice carrying the *Flp1* recombinase gene under control of the *Actb* promoter (005703, Jackson Laboratory, Bar Harbor ME). Resulting mice carrying the *Sar1b*^*fl*^ allele were then bred with mice carrying an *EIIa-Cre* transgene (003724, Jackson Laboratory, Bar Harbor ME) to generate the *Sar1b* null allele (*Sar1b*^*-*^) (**Figure 3A**). *Sar1b*^*fl/+*^ and *Sar1b*^*+/-*^ mice were maintained by continuous backcrossing to C57BL/6J mice.

### Generation of *Sar1b*^*fl/+*^ *Alb-Cre*^*+*^ mice

To generate *Sar1b*^*fl/+*^ *Alb-Cre*^*+*^ mice, the *Sar1b*^*fl/+*^ mice generated above were crossed with mice carrying an *Alb-Cre* transgene (003574, Jackson Laboratory, Bar Harbor ME) (14). *Sar1b*^*fl/+*^ *Alb-Cre*^*+*^ mice were maintained by continuous backcrossing to C57BL/6J mice. *Sar1b*^*fl/fl*^ *Alb-Cre*^*+*^ mice were obtained by crossing *Sar1b*^*fl/+*^ *Alb-Cre*^*+*^ mice with *Sar1b*^*fl/+*^ or *Sar1b*^*fl/fl*^ mice.

### Generation of *Sar1b*^*+/a*^ mice

Mice carrying the *Sar1b*^*a*^ allele, in which the mouse *Sar1a* coding sequence is inserted into the *Sar1b* locus immediately downstream of the ATG codon in exon 2 of the gene, were generated by Biocytogen Boston Corp. (Wakefield MA) using CRISPR/Cas9 (**Figure 5A**). Cas9 and an sgRNA targeting exon 2 of *Sar1b* along with the DNA repair template were injected into C57BL/6N zygotes. Founder mice with confirmed germ-line transmission were bred to C57BL/6N mice to obtain F1 (*Sar1b*^*+/a*^) mice. *Sar1b*^*+/a*^ mice were backcrossed to C57BL/6J mice for at least 3 generations before experimentation. *Sar1b*^*+/a*^ mice were maintained by continuous backcrossing to C57BL/6J mice.

### Mouse genotyping

Tail clips were obtained from 2-3 weeks old mice for genomic DNA isolation and genotyping, as previously described (10). PCR was performed using Go-Taq Green Master Mix (Promega, Madison, WI) and resulting products were resolved by 3% agarose gel electrophoresis. All primers used for genotyping are listed in **Table S2**.

### Timed mating and embryo collection

Timed matings were performed by intercrossing heterozygous mice (*Sar1a*^*gt/+*^ or *Sar1b*^*+/-*^). Embryos were collected at E10.5, E11.0, E11.5, or E18.5 for genotyping and histologic analyses. Genomic DNA was isolated from the embryonic yolk sacs of E10.5-E11.5 embryos or tail clips of E18.5 embryos and genotyped as described above. Embryos were immediately fixed in Z-fix solution (Anatech Ltd, Battle Creek MI) for subsequent histologic analyses.

### Blood and tissue collection

Blood and tissues were collected and sera separated as previously described (35). Tissues were immediately fixed in Z-fix solution or frozen in liquid nitrogen. Sera and tissues were stored at -80°C until experimentation.

### Histologic analyses

Tissue processing, embedding, sectioning, hematoxylin and eosin (H&E) staining were performed at the University of Michigan In-Vivo Animal Core (IVAC). Slides were reviewed by the investigators and a veterinarian pathologist blinded to genotype.

### Cholesterol assays

Total cholesterol in sera was determined using a colorimetric assay (SB-1010-225, Fisher Scientific, Hampton NH) according to the manufacturer’s instructions.

### Quantitative PCR (qPCR) assays

Total RNA was isolated from tissues and converted to cDNA as previously described (35). Primers specific for each target (**Table S3**) were designed using NCBI Primer-BLAST. Quantitative PCR reactions were performed using Power SYBR Green PCR Master Mix (4367659, Invitrogen, Waltham MA). *Gapdh, B2m, Rpl37*, and *Rpl38* were used as housekeeping gene controls. Normalized transcript abundance was calculated by the 2^-ΔΔCt^ method using the gene with the lowest standard deviation across all samples as the endogenous control.

### Adenovirus mediated expression of SAR1A or SAR1B

Recombinant adenoviruses expressing mouse SAR1A or SAR1B were constructed using pAdTrack-CMV and the AdEasy adenoviral vector system (Agilent Technologies, Lexington MA). Adenoviruses were amplified in Ad293 cells and purified using CsCl gradient ultracentrifugation. Mice were transduced with Ad-GFP, Ad-SAR1A, or Ad-SAR1B via tail vein injection at 0.1 OD per mouse (low dose) or 0.3 OD per mouse (high dose), as described previously (36). Plasma cholesterol concentrations were measured prior to and 3 or 7 days after transduction as described above.

## Acknowledgments

This work was supported by NIH grants R35HL135793 (DG), R01HL148333 (RK), R01HL157062 (RK), K08HL148552 (BTE), and R00GM141268 (PSA). VTT was supported by fellowships from the American Heart Association (20PRE35210706) and the University of Michigan Horace H. Rackham School of Graduate Studies.

---

**Table S1:**
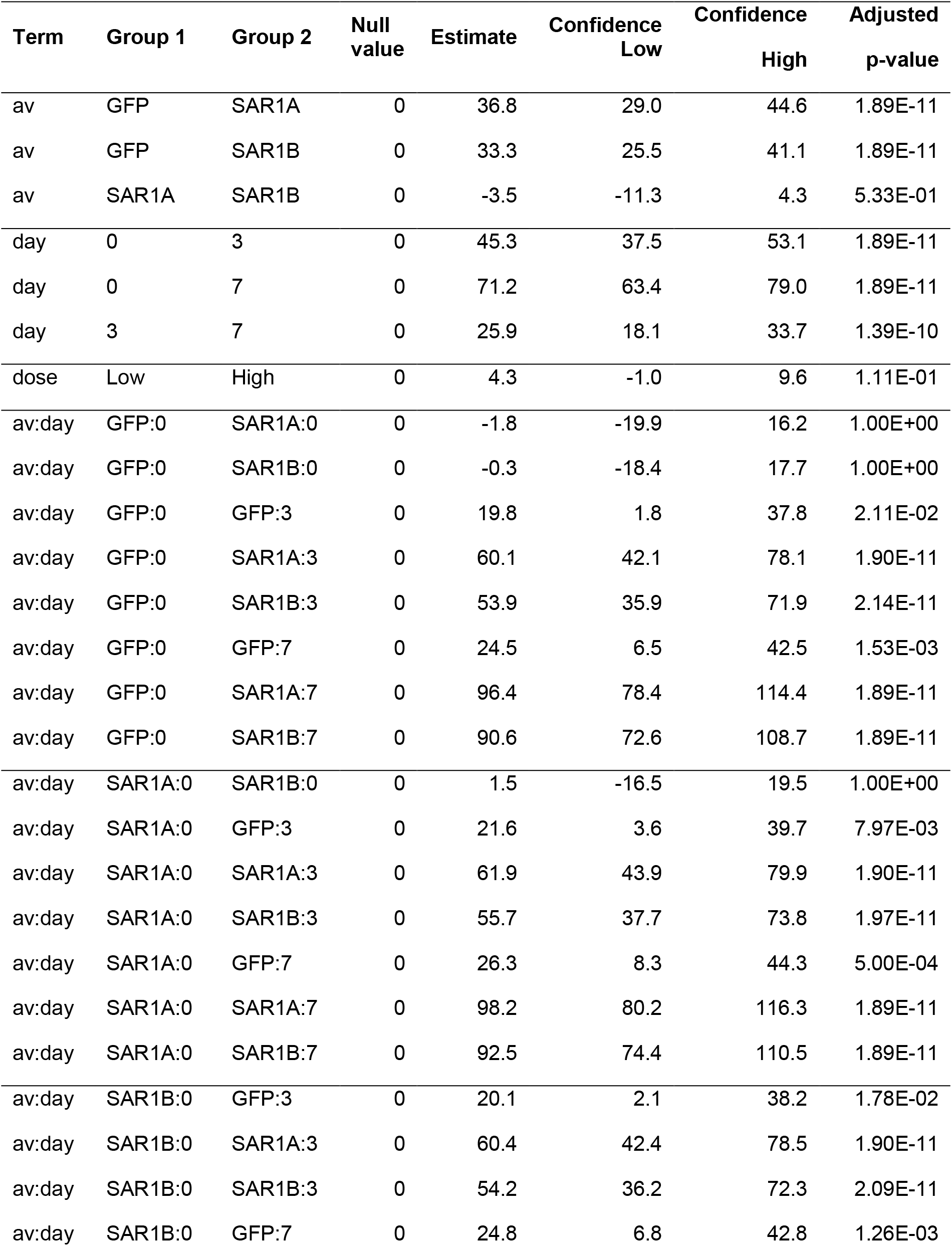
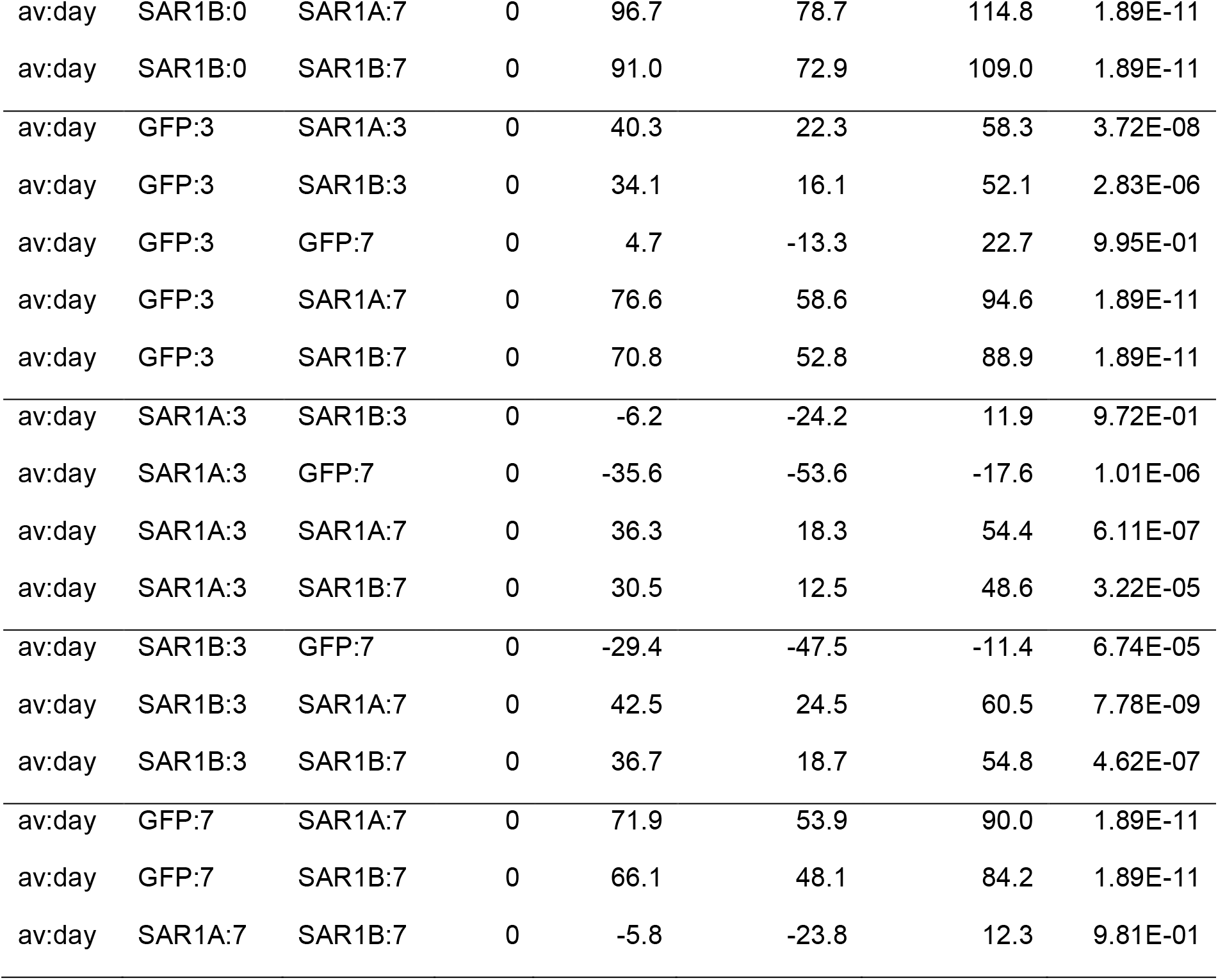
Tukey’s post-hoc test for comparison between AV type, time, and dose.

**Table S2:**
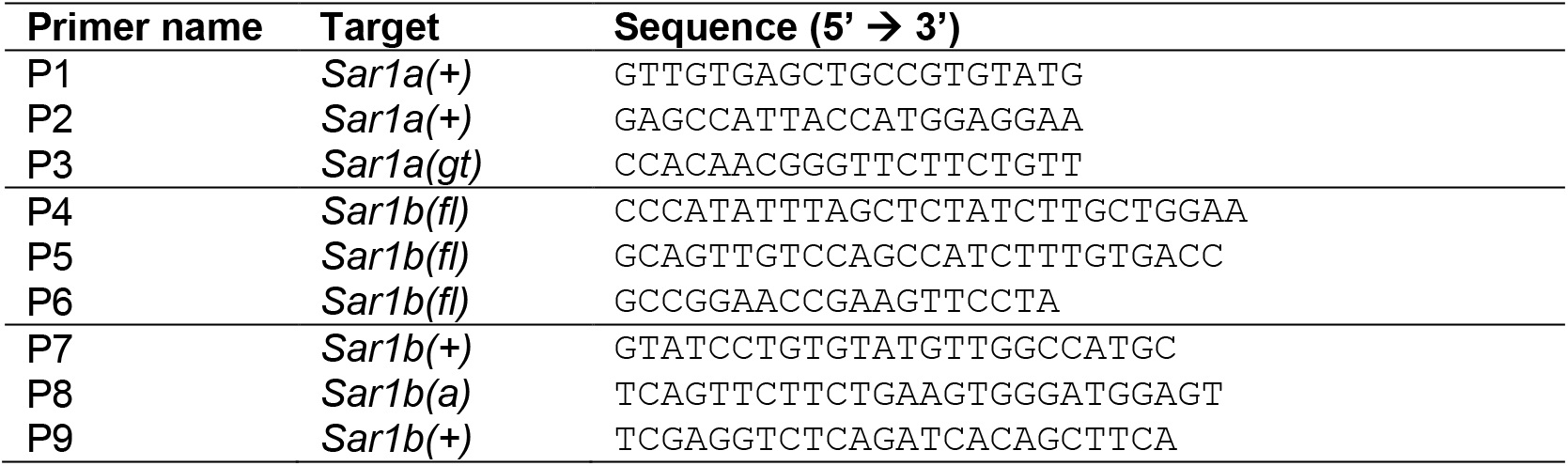
Genotyping primer sequences.

**Table S3:**
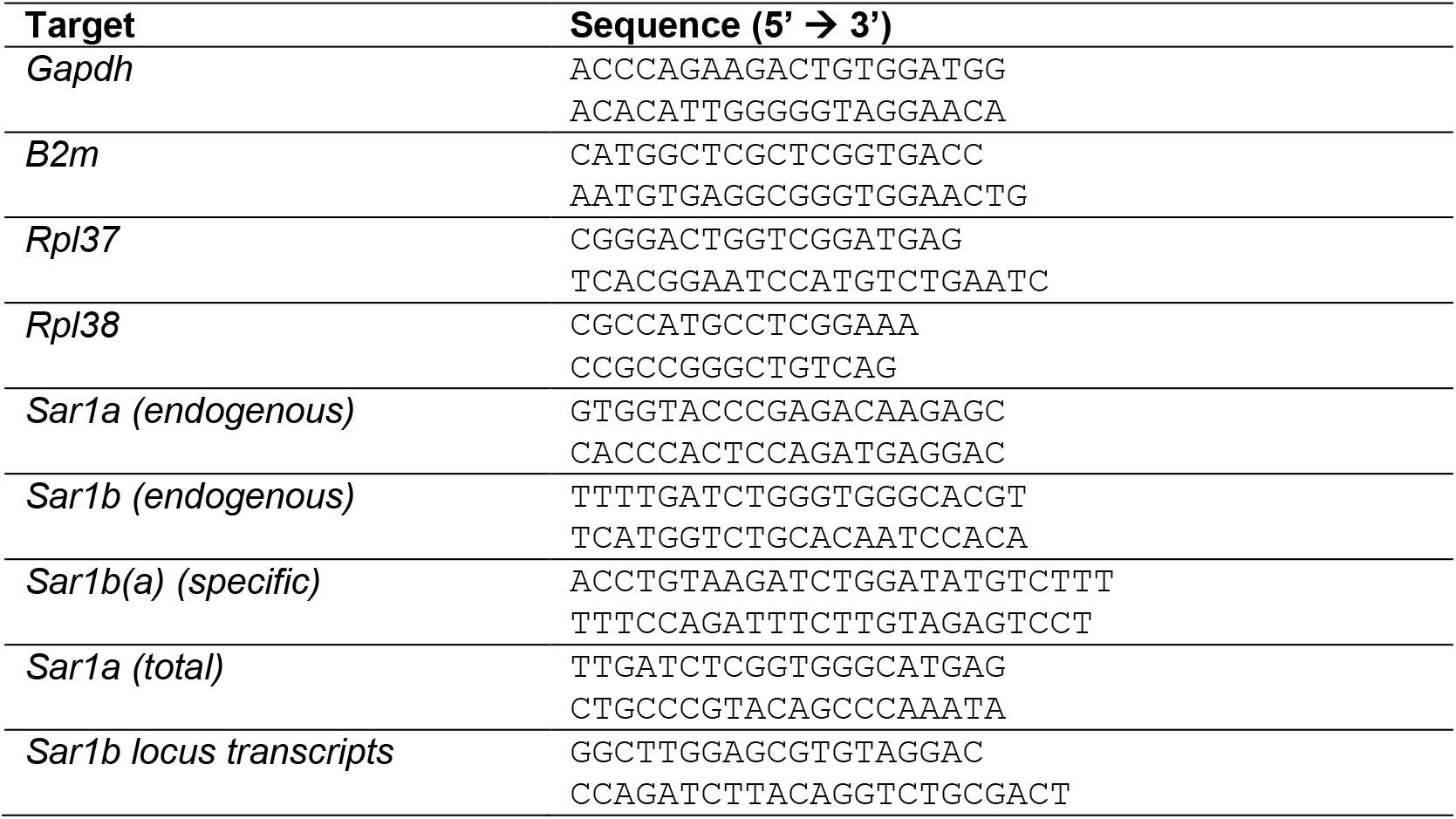
Primers used for qPCR assays.

